# Crosslinking Proteomics Indicates Effects of Simvastatin on the TLR2 interactome and Reveals ACTR1A as a Novel Regulator of the TLR2 Signal Cascade

**DOI:** 10.1101/530402

**Authors:** Abu Hena Mostafa Kamal, Jim J. Aloor, Michael B. Fessler, Saiful M. Chowdhury

**Affiliations:** Department of Chemistry and Biochemistry, University of Texas at Arlington, Texas 76019, USA; Immunity, Inflammation and Disease Laboratory, National Institute of Environmental Health Sciences, National Institutes of Health, Research Triangle Park, NC 27709, USA

**Keywords:** Alpha-centractin, arp-1, Dynactin, Pam3CSK4, Cross-linkers, Toll-like receptor 2, Simvastatin, Affinity Proteomics, Mass Spectrometry, Immunoprecipitation

## Abstract

Toll-like receptor 2 (TLR2) is a pattern recognition receptor that, upon ligation by microbial molecules, interacts with other proteins to initiate pro-inflammatory responses by the cell. Statins (hydroxymethylglutaryl coenzyme A reductase inhibitors), drugs widely prescribed to reduce hypercholesterolemia, are reported to have both pro- and anti-inflammatory effects upon cells. Some of these responses are presumed to be driven by effects on signaling proteins at the plasma membrane, but the underlying mechanisms remain obscure. We reasoned that profiling the effect of statins on the repertoire of TLR2-interacting proteins might provide novel insights into the mechanisms by which statins impact inflammation. In order to study the TLR2 interactome, we designed a co-immunoprecipitation (IP)-based cross-linking proteomics study. A hemagglutinin (HA)-tagged-TLR2 transfected HEK293 cell line was utilized to precipitate the TLR2 interactome upon cell exposure to the TLR2 agonist Pam3CSK4 and simvastatin, singly and in combination. To stabilize protein interactors, we utilized two different chemical cross-linkers with different spacer chain lengths. Proteomic analysis revealed important combinatorial effects of simvastatin and Pam3CSK4 on the TLR2 interactome. After stringent data filtering, we identified alpha-centractin (ACTR1A), an actin-related protein and subunit of the dynactin complex, as a potential interactor of TLR2. The interaction was validated using biochemical methods. RNA interference studies revealed an important role for ACTR1A in induction of pro-inflammatory cytokines. Taken together, we report that statins remodel the TLR2 interactome, and we identify ACTR1A, a part of the dynactin complex, as a novel regulator of TLR2-mediated immune signaling pathways.

## Introduction

Protein interactions have an important role in biological and cellular systems, including gene expression, signaling, and immune responses. The challenges associated with identifying specific protein-interacting partners in complex biological samples (1–3) have led to the development of several methodological approaches. Co-immunoprecipitation (IP)-based identification of protein interactions is a gold standard technique for defining protein complexes in native biological systems (4). In this method, a protein of interest is subjected to affinity- or antibody-based purifications along with its interacting partners. Optimization of wash conditions that remove non-specific interactions but preserve transient and weak interactions is a major challenge that renders this method most amenable to identifying stable protein-protein interactions. In order to improve co-IP proteomics, protein cross-linking methods that covalently attach proximal protein binding partners have recently been employed (5, 6). Cross-linking theoretically captures transient and weak protein interactions, permitting the subsequent use of strong denaturing washing conditions that preserve specificity. A further advantage of cross-linking methods is that interactions can be defined either through identifying the proteins or in some cases through specifically examining cross-linked peptides. While domain-specific cross-linking data analysis is hindered due to the complexity of bioinformatics software, several software packages are currently available for specific cross-linkers. Nonetheless, because confident protein identification is still very challenging for large-scale data sets, identifying the interaction of cross-linked proteins by examining unmodified peptides has become a very popular approach.

The Toll-Like Receptors (TLRs) are a family are type I transmembrane proteins of the innate immune system that trigger a stereotypical pro-inflammatory cytokine induction response upon ligation. The 10 TLRs of the human innate immune system are localized to either the plasma membrane (TLR1, 2, 4, 5, 6) or endosome (TLR3, 7, 8, 9), and are activated by signature molecular patterns present in microbes (7–11). Of the TLRs, TLR2, activated by lipoteichoic acid, synthetic lipopeptides (Pam3CSK4 [P3C]), and glycans from Gram-positive bacteria, Gram-negative bacteria, and mycobacteria (12, 13), plays a pivotal role in the early innate immune response to microbial infections through triggering a signaling cascade that leads to the activation of the pro-inflammatory transcription factor nuclear factor-κB (8, 13, 14). Moreover, TLR2-dependent signaling contributes to the pathogenesis of a wide variety of diseases, such as antiphospholipid syndrome, sepsis, tuberculosis, chronic obstructive pulmonary, cytomegalovirus infection, rheumatic heart disease, cerebral injury, cutaneous leishmaniasis, atherosclerosis and leptospirosis (16–22). Given this, an improved understanding of TLR2-dependent signaling and how it is altered by immunomodulatory medications may provide novel insights relevant to human disease.

Statins, inhibitors of 3-hydroxy-3-methylglutaryl-CoA (HMG-CoA) reductase, are widely prescribed to reduce serum cholesterol in hyperlipidemic patients. Recently, statins have been shown to have additional immunomodulatory activities that are relevant to the pathogenesis of cardiovascular and other diseases (23). For example, statins suppress maturation of human monocyte-derived dendritic cells (24), and have been shown to ameliorate inflammation in a wide range of animal models of immunological disorders such as autoimmune encephalomyelitis (25, 26), sepsis (27), and graft arterial disease (GAD). Paradoxically, in other settings, proinflammatory effects of statins have also been identified on several cell types, including endothelial cells, peripheral blood mononuclear cells, and dendritic cells (28). The mechanisms that determine the pro-vs. anti-inflammatory actions of statins in different settings remain poorly understood, but in many cases have been proposed to stem from indirect effects on membrane proteins (e.g., receptors).

We reasoned that P3C activation of TLR2 would serve as a well-defined model system amenable to unbiased proteomic-based profiling of the receptor-proximal effects of statins upon inflammatory signaling. In order to study the combinatorial effects of P3C and statins on the TLR2 protein interactome, we designed a cross-linking-based co-IP MS strategy. HEK293 cells stably expressing HA-tagged TLR2 were used to pull down TLR2 along with its interactors following crosslinker treatment. Given that smaller cross-linking agents may miss covalent attachment of surface receptors and cytosolic proteins near the membrane, we utilized two cross-linker agents with different spacer-chain lengths, our recently developed a dual cleavable cross-linker (DUCCT; spacer chain distance ~16.3 Å) (29) and a commercial cross-linker BS3 (spacer chain distance, 11.4 Å). After crosslinking and affinity pulldown, proteins were separated by SDS-PAGE, trypsin-digested, and the resultant peptides were analyzed by liquid chromatography-tandem mass spectrometry (LC-MS/MS) followed by data filtering.

Two proteins, alpha-centractin (ACTR1A) and myristoylated alanine-rich C kinase substrate-like protein 1 (MARCKSL1), were identified as novel interactors of TLR2 exclusively in statin-treated cells under DUCCT cross-linker treatment. We followed this discovery up with biochemical validation studies. We show that ACTR1A has important modulatory actions on the TLR2 pro-inflammatory signaling cascade. Taken together, these data identify for the first time that ACTRA1 is a statin-responsive protein that serves to modulate TLR2-mediated signaling. Given the prevalence of statin use in human populations, these mechanistic studies may have important translational implications.

### Experimental procedures

#### HA-TLR2-MD2-CD14-293 cell line

A plasmid expressing the HA-tagged human TLR2 gene (Catalog # puno-htlr2ha, Invivogen, San Diego, CA) was transiently transfected into HEK293-hMD2-CD14 cells (Invivogen) using Lipofectamine 2000. Standard antibiotic selection procedures using Blasticidin S hydrochloride (Invivogen) were utilized together with western-blotting and immunostaining verification, to generate stable cell lines that strongly expressed (driven by a composite promoter hEF1/HTLV) the hTLR2-HA protein. In order to maintain selective pressure, the cell line was grown and maintained in DMEM containing 10% FBS, 100 U/ml penicillin, and 100μg/ml streptomycin, supplemented with Blasticidin 10μg/ml and Hygromycin 50μg/ml.

### Cell culture and protein preparation

Hemagglutinin (HA) tagged-TLR2-MD2-CD14-human embryonic kidney (HEK)293 cells were maintained in DMEM supplemented with 10% fetal bovine serum, 1% penicillin/streptomycin in a humidified atmosphere of 5% CO_2_, and antibiotics (50 μg/ml hygromycin and 10 μg/ml blasticidin). Cells were treated with 10 μM simvastatin (Sigma) for 24 hr, then stimulated with 1 μg/ml Pam3CSK4 (P3C; InvivoGen) for 24 hr in fresh medium. The cells were then treated with Dual Cleavable Cross-linking Technology (DUCCT) (29) or commercial bissulfosuccinimidyl suberate (BS3) cross-linker (XL), added at a final concentration of 1μmol/ml for 30 min, followed by quenching the reaction with 50 mM Tris-HCl, pH 8.0. For IP-pull down for proteomics, the cells were lysed with immunoprecipitation (IP)-lysis buffer supplemented with protease inhibitors at 4°C for 15 mins, then sonicated for another 15 mins. Finally, the suspended cells were kept at 4°C for 30 mins, then centrifuged (20,000×g, 4°C, 30 min). The supernatant was collected for measuring the protein concentration with a BCA protein assay kit, using bovine serum albumin as a standard.

### Separating the TLR2-interacting partners using immunoprecipitation

Anti-HA magnetic beads (Thermo Scientific, MA) were washed with 0.05% TBS-T buffer and gently vortexed. Suspended magnetic beads were collected using magnetic stand for 5 min at room temperature (RT). HA-tagged protein samples were mixed into the pre-washed beads and gently rotated at 4°C overnight. The beads were then collected with a magnetic stand and washed with TBS-T buffer and ultrapure-water twice, followed by elution in Laemmli buffer (95°C, 5 min). After centrifugation, the reduced samples were loaded onto SDS-PAGE gels (12%) for separation, followed by staining with Sypro Ruby in the dark for 12 hr (Fig. S1).

For reverse co-immunoprecipitation (IP), protein samples were mixed with 5-10 μg anti-ACTR1A or -MARCKSL1 antibody, after volume adjustment to 500 μl with IP lysis buffer. The samples were incubated for overnight at 4°C with continuous mixing, then exposed to washed protein G magnetic beads (Thermo Scientific, MA) and incubated overnight at 4°C with continuous mixing. Beads were collected using a magnetic stand, washed with washing buffer and ultra-pure water, then eluted in Laemmli buffer (95°C, 5 min). The protein eluent was then separated by SDS-PAGE for immunoblotting.

### In-gel digestion and mass analysis (nano-LC-MS/MS)

SDS-PAGE gel bands were excised, squeezed with acetonitrile, and dried at room temperature. Proteins were then reduced and alkylated and digested with trypsin (porcine) (MS Grade) at 37°C for overnight (30). Formic acid to pH < 3 was added to the resulting peptides, followed by drying by speed vacuum, and then dissolution in 0.1% formic acid. Finally, the peptides were centrifuged at 20,000 × g for 30 min at 4°C.

Digested peptides were analyzed by nano LC-MS/MS using a Velos Pro Dual-Pressure Linear Ion Trap Mass Spectrometer (ThermoFisher Scientific, MA) coupled to an UltiMate 3000 UHPLC (ThermoFisher Scientific, MA). Peptides were loaded onto the analytical column and separated by reverse-phase chromatography using a 15-cm column (Acclaim PepMap RSLC) with an inner diameter of 75 μm and packed with 2 μm C_18_ particles (Thermo Fisher Scientific, MA). The peptide samples were eluted from the nano column with multi-step gradients of 4-90% solvent B (A: 0.1% formic acid in water; B: 95% acetonitrile and 0.1% formic acid in water) over 70 min with a flow rate of 300 nL/min with a total run time of 90 min. The mass spectrometer was operated in positive ionization mode with nano spray voltage set at 2.50-3.00 kV and source temperature at 275°C. The three precursor ions with the most intense signal in a full MS scan were consecutively isolated and fragmented to acquire their corresponding MS2 scans. Full MS scans were performed with 1 micro scan at resolution of 3000, and a mass range of m/z 350-1500. Normalized collision energy (NCE) was set at 35%. Fragment ion spectra produced via high-energy collision-induced dissociation (CID) was acquired in the Linear Ion Trap with a resolution of 0.05 FWHM (full-width half maximum) with an Ultra Zoom-Scan between *m*/*z* 50-2000. A maximum injection volume of 5 μl was used during data acquisition with partial injection mode. The mass spectrometer was controlled in a data-dependent mode that toggled automatically between MS and MS/MS acquisition. MS/MS data acquisition and processing were performed by Xcalibur™ software, ver. 2.2 (ThermoFisher Scientific, MA).

### Database search

Proteins were identified through Proteome Discoverer software (ver. 2.1, Thermo Fisher Scientific) using UniProt human (*Homo sapiens*) protein sequence database (120672 sequences, and 44548111 residues). The reviewed protein sequences of human were downloaded from UniProt protein database (www.uniprot.org) on August 12, 2016. The considerations in SEQUEST searches for normal peptides were used with carbamidomethylation of cysteine as the static modification and oxidation of methionine as the dynamic modification. Trypsin was indicated as the proteolytic enzyme with two missed cleavages. Peptide and fragment mass tolerance were set at ± 1.6 and 0.6 Da and precursor mass range of 350-3500 Da, and peptide charges were set excluding +1 charge state. SEQUEST results were filtered with the target PSM validator to improve the sensitivity and accuracy of the peptide identification. Using a decoy search strategy, target false discovery rates for peptide identification of all searches were < 1% with at least two peptides per protein, a maximum of two missed cleavage, and the results were strictly filtered by ΔCn (< 0.01), Xcorr (≥ 1.5) for peptides, and peptide spectral matches (PSMs) with high confidence, that is, with *q*-value of ≤ 0.05. Proteins quantifications were conducted using the total spectrum count of identified proteins. Additional criteria were applied to increase confidence that PSMs must be present in all three biological replicates samples. Normalization of identified PSMs among LC-MS/MS runs was done by dividing individual PSMs of proteins with total PSMs and average of % PSM count was utilized for calculating fold changes for different treatment conditions (30, 31). For contrasting relative intensities of proteins between control, P3C, statin-P3C, and statin groups, samples were evaluated using cumulative confident normalized PSMs value.

### Gene ontology and protein interaction analysis

Protein-encoding genes were functionally categorized using gene ontology systems by PANTHER classification system-based biological processes, molecular activities, and cellular components (32). Protein abundances were visualized as a heat map. The cluster was generated by MeV software (ver. 4.9; http://www.tm4.org/) (33). The proteomic data set, which included UniProt identifiers and fold changes of total identified protein, was submitted into Ingenuity Pathway Analysis (IPA) for core analysis (Ingenuity Systems, Redwood City, CA). The matched proteins with submitted dataset in Ingenuity Knowledge Base generated TLR2 protein interaction networks according to biological as well as molecular functions. The core analysis was performed with the settings of indirect and direct relationships between molecules based on experimentally observed data, and data sources were considered in human databases in the Ingenuity Knowledge Base (34). For generating the protein interaction networks in proteins exclusively identified upon treatment with DUCCT- and BS3-XLs, identified protein-coding genes were submitted into the Cytoscape ver. 3.6.1 according to affinity purification-mass spectrometry protein network analysis methods (35).

### Immunoblotting

For immunoblotting, cells were washed with 1x PBS twice and then lysed with RIPA buffer (same as protein preparation). Protein samples were prepared in 2X Laemmli buffer and were heated for 5 min at 95°C. Proteins were separated on a 12% polyacrylamide gel. The proteins were transferred to a 0.45 μm nitrocellulose membrane for 1.5 hr at 100 V. The nitrocellulose membrane was then blocked in skim milk (5%) in TBST buffer for 2 hr at room temperature (RT) and incubated with primary antibodies against ACTR1A (ab203833; Abcam), MARCKSL1 (ab184546; Abcam), or TLR2 (ab191458; Abcam) in bovine serum albumin (5%) at 4°C for overnight. Goat anti-rabbit IgG secondary antibody conjugated to HRP (Abcam) was then used for 2 hr at RT. β-actin (ab8227; Abcam), and GAPDH (ab9485; Abcam) antibodies were used as loading controls. The targeted protein bands were visualized using clarity western enhanced chemiluminescent substrate (BioRad).

### Immunocytochemistry

Cells were grown on 1M HCL-treated glass slides, and then fixed with chilled methanol for 5 min at RT. Cells were subsequently permeabilized with 0.1% Triton X-100 in 1x PBS for 10 min, blocked with bovine serum albumin and glycine in 1x PBS for 30 min at RT in the dark, and then incubated with anti-ACTR1A (ab11009, Abcam) or anti-TLR2 (PA5-20020, ThermoFisher scientific, IL, USA) antibodies at 4°C overnight in the dark. They were then incubated with secondary antibody (goat anti-rabbit IgG H&L, Alexa Flour 488, ab15007, Abcam) (2 hr, dark, room temperature), and imaged with a Leica DMi8 confocal microscope (Leica, IL, USA). The images were analyzed using Lax X (Leica, IL, USA). DAPI was used for nuclear staining.

### Transfection with ACTR1A siRNA and qRT-PCR

ACTR1A-targeted small interfering RNAs (siRNA; SR306823 for human) and nonsense siRNA were purchased from OriGene (OriGene, MD, USA). For transfections, HEK293 cells were seeded in 6-well plates with DMEM medium supplemented with 10% FBS, 1% penicillin/streptomycin, and selective antibiotics (see cell culture methods). After 50~70% confluence, cells were transfected according to the manufacturer’s instructions. After 48 hr, cells were treated with statin for 24 hr and P3C for an additional 24 hr. Following this, the cells were collected for mRNA analysis. Total RNA was extracted by TRIzol^®^ (Invitrogen) (30). First-strand cDNA synthesis was performed using one-step cDNA synthesis kit (Origene, MD, USA). Real-time PCR was performed on the CFX96 real-time system (Bio-Rad) using the *SsoAdvanced*™ Universal SYBR^®^ *Green* Supermix (Bio-Rad). Each assay was performed in triplicate, and the mean value was used to calculate the mRNA expression for the gene of interest and the housekeeping reference gene (GAPDH). The abundance of the gene of interest in each sample was normalized to that of the reference control using the comparative (2^–ΔCT) method (36). Sequences of the primers are provided in the supplementary information (Table S1).

### Statistical analysis

The quantitative analysis of proteins as PSMs was performed using built-in-statistical packages in Proteome Discoverer (Ver. 2.1). Results were considered statistically significant if *q* ≤ 0.05. Scatter plots and pairwise correlation matrices were generated using the R package, where results were considered if correlation coefficient (*R*^2^) was > 0.80. The data are depicted in the graphs as mean ± standard error (SEM). Statistical significance was determined using one-way ANOVA with *P* ≤ 0.05 considered as significant. GraphPad Prism version 6 was used (GraphPad Software, Inc).

## Results

### Identification of TLR2-interacting proteins

To identify the effect of P3C and statins on the TLR2 interactome, we performed co-IP proteomics on HA-TLR2-MD2-CD14-HEK293 cells from four exposure conditions (control; P3C; statin; statin-P3C) following post-exposure treatment with DUCCT or BS3 crosslinker (Fig. 1). Control samples untreated with crosslinker were also analyzed. After pulldown with anti-HA magnetic beads, precipitated proteins were separated by SDS-PAGE (Fig. S1) and the resulting gel bands were digested using trypsin and then analyzed by nano-LC-MS/MS and database searching (UniProt). Peptides were quantified using Peptide Spectrum Matches (PSMs). Correlation matrix comparisons among three biological replicates are shown in Fig. S2. Pairwise correlation coefficients among the biological replicates showed high correlation with a *R*^2^ value of >0.80.

**Figure 1.**
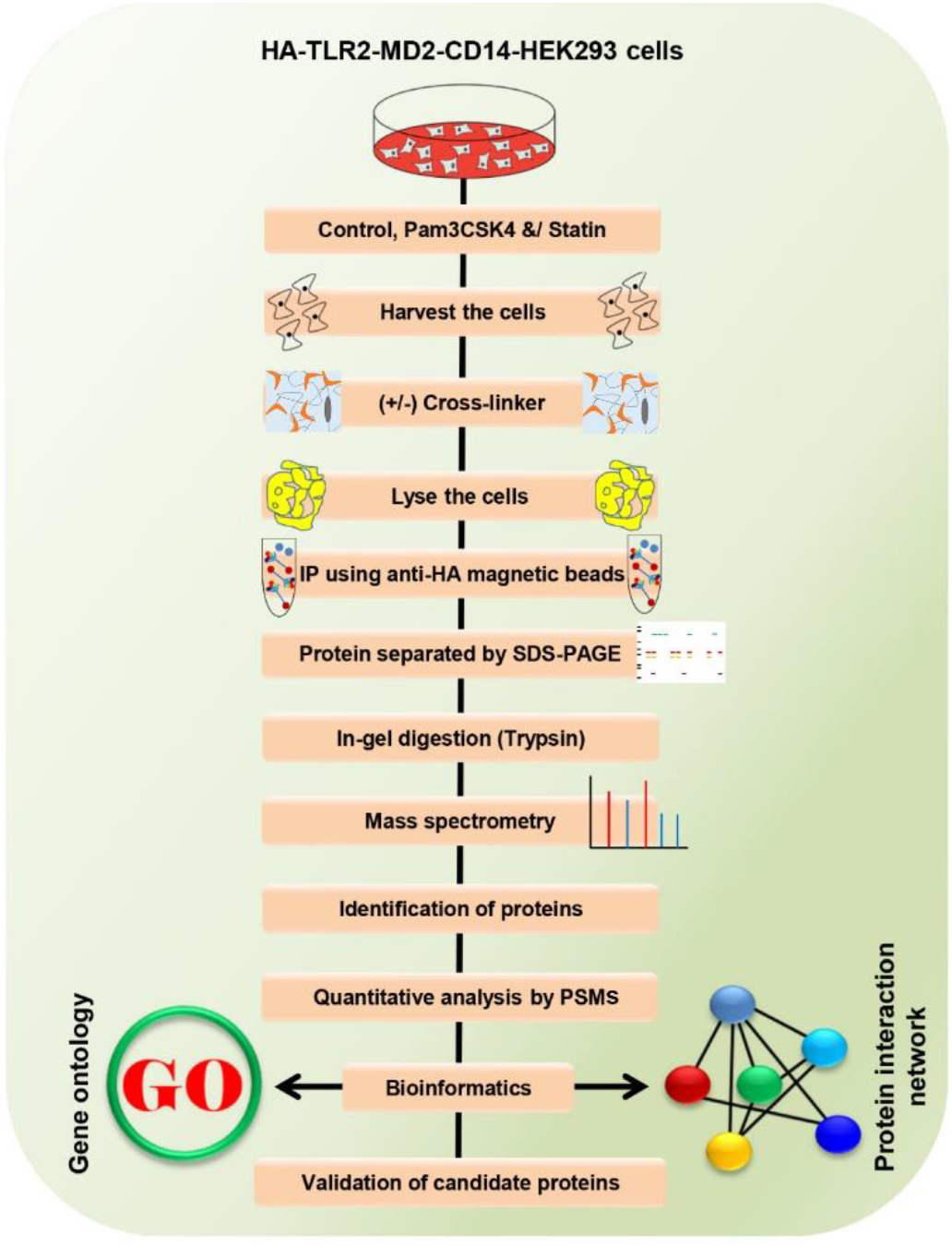
The experimental procedure of the IP-crosslinker-MS-based proteomic analysis. HEK293 cells were treated with statin and Pam3CSK4 along with cross-linkers, as depicted. Pull-down samples were separated by SDS-PAGE and analyzed by nano-LC-MS/MS, then quantitative analysis was performed by PSMs. Different molecular techniques were used for characterizing the candidate proteins during immune responses.

Overall, 1153 proteins were identified and quantified across all conditions. The data set was filtered using two unique peptides per protein and a false discovery rate of 1%. Detailed information about the identification of proteins and peptides is shown in Table S2-S4. First, we examined proteins that were identified across the four cell exposure conditions, but in the absence of crosslinker treatment. Out of 803 proteins, 249 proteins were common to all four exposures, whereas 249 proteins were exclusive to a single exposure (16 in control, 78 in P3C, 149 in statin-P3C, and 6 in statin) (Fig. 2A). We next evaluated the impact of the two cross-linkers on protein discovery in the setting of HA-TLR2 pulldowns from P3C, statin, and P3C-statin-treated cells. In samples treated with DUCCT in combination with P3C, 220 proteins were commonly shared across control, P3C, P3C-DUCCT and DUCCT conditions, whereas 288 proteins were exclusively identified in individual conditions. Consistent with improved protein recovery with DUCCT, 16 more proteins were identified in P3C-DUCCT samples (total, 589 proteins) than in P3C-stimulated samples without DUCCT (total, 605 proteins) (Fig. 2B and Table S2). Of these, 147 proteins were exclusive to P3C-DUCCT (i.e., not detected under P3C, control, or DUCCT conditions) (Fig 2B). Regarding statin-P3C-cotreated samples, 167 proteins were identified exclusively in statin-P3C samples, whereas, 28 proteins were exclusive to statin-P3C-DUCCT samples (Fig. 2C). In comparison of statin and statin-DUCCT treated samples (Fig. 2D), 15 and 221 proteins were exclusively identified, respectively.

**Figure 2.**
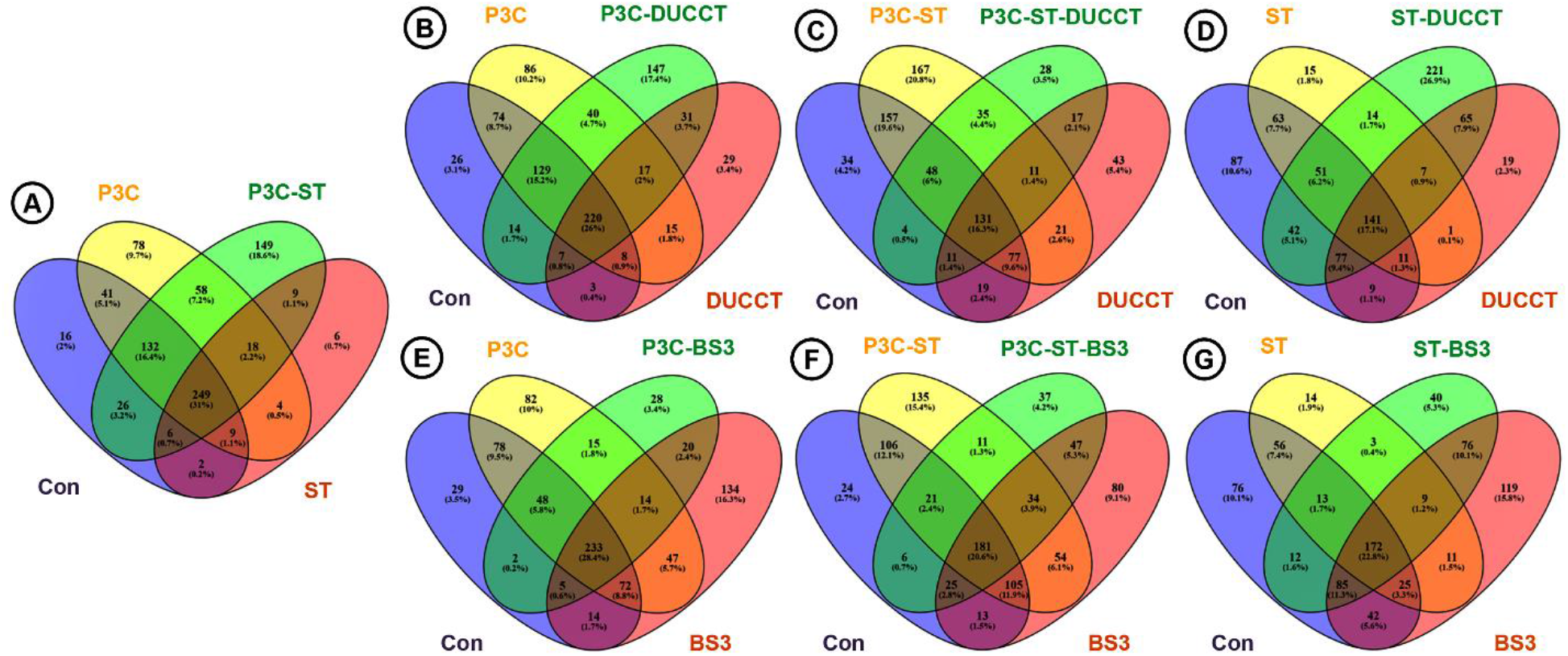
Venn diagrams showing distributions of identified proteins across cell treatment conditions. The diagrams show the distributions of the total identified proteins in HEK293 cells upon treatment with statin (ST) and Pam3CSK4 (P3C; A), and in Pam3CSK4-, Pam3CSK4-statin, and statin-treated HEK293 cells along with the DUCCT crosslinking (B-D) or BS3 crosslinking (E-G).

Distinct effects on the TLR2 interactome were noted with BS3 cross-linker. Contrary to the increase in protein recovery in P3C-treated cells enforced by DUCCT, BS3 treatment led to 224 fewer proteins identified under P3C treatment conditions (Fig. 2E). Because of this, remarkably, 240 more proteins were identified in DUCCT-treated P3C samples than in BS3-treated P3C samples (compare Figs. 2B and 2E). Similarly, 285 fewer proteins were identified in statin-P3C-BS3 samples than in statin-P3C samples (Fig. 2F). In this case, however, 77 more proteins were identified in BS3 samples compared to DUCCT samples after statin-P3C (Fig. 2C and 2F). Finally, in the case of statin-treated cells, BS3 led to identification of 107 more proteins (Fig. 2G). Due to a more marked improvement in protein recovery with DUCCT, 208 more proteins were identified in DUCCT-treated samples than in BS3-treated samples following statin exposure (Fig. 2D and 2G). Taken together, we conclude that, overall, compared to BS3, the DUCCT crosslinker led to improved recovery of the TLR2 interactome.

Visualization by heat map of the relative expression (normalized PSMs) of TLR2-interacting proteins (N=803) suggests that treatment with P3C, statins, and P3C-statins induce distinct biological states of the cells (Fig. 3A). Different proteomic expression patterns were evident among 1082 proteins identified under DUCCT treatment (Fig. 3B) and the 975 protein intensities identified under BS3 treatment (Fig. 3C), confirming that the two cross-linkers enforce a significant effect on the detected TLR2 interactome.

**Figure 3.**
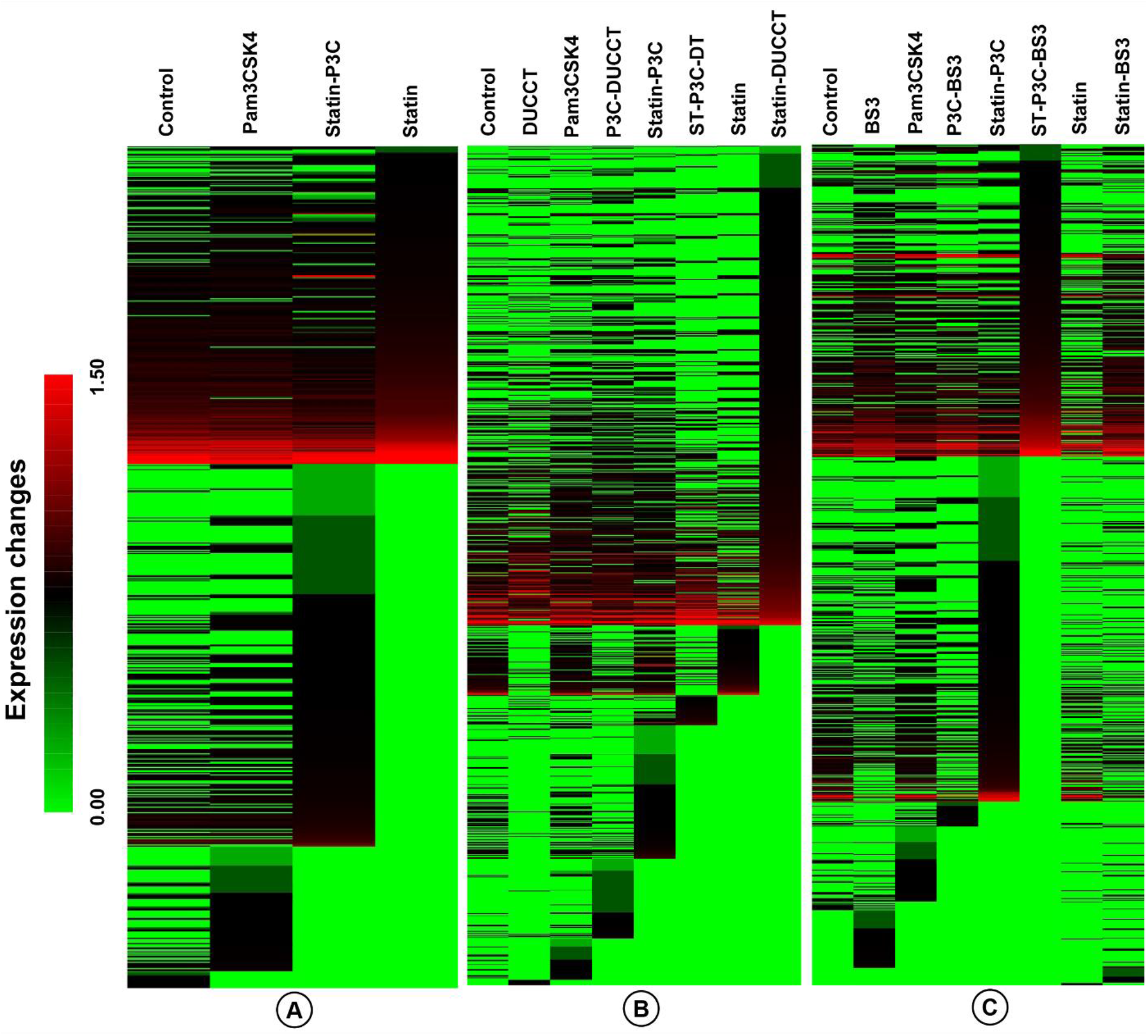
Heatmap showing the relative expression levels of proteins across cell treatment conditions. Proteins were differentially expressed in HEK293 cells upon the treatment of statin (ST) and Pam3CSK4 (P3C; A), along with cross-linkers-DUCCT (DT) (B) and BS3 (C).

All 803 proteins identified in the absence of cross-linker treatment were categorized based on gene ontology using PANTHER gene classification systems (32). Overall, 8, 11, and 8 ontology pathways were represented within the ontology categories of cellular components, biological processes, and molecular functions, respectively. Several ontology pathways were particularly enriched by P3C and statin treatment. These are ‘cell part’ and ‘organelle’ under the ‘cellular components’ category, ‘metabolic process’ and ‘cellular process’ under the ‘biological processes’ category, and ‘catalytic activity’ and ‘binding’ under the ‘molecular functions’ category (Fig. 4).

**Figure 4.**
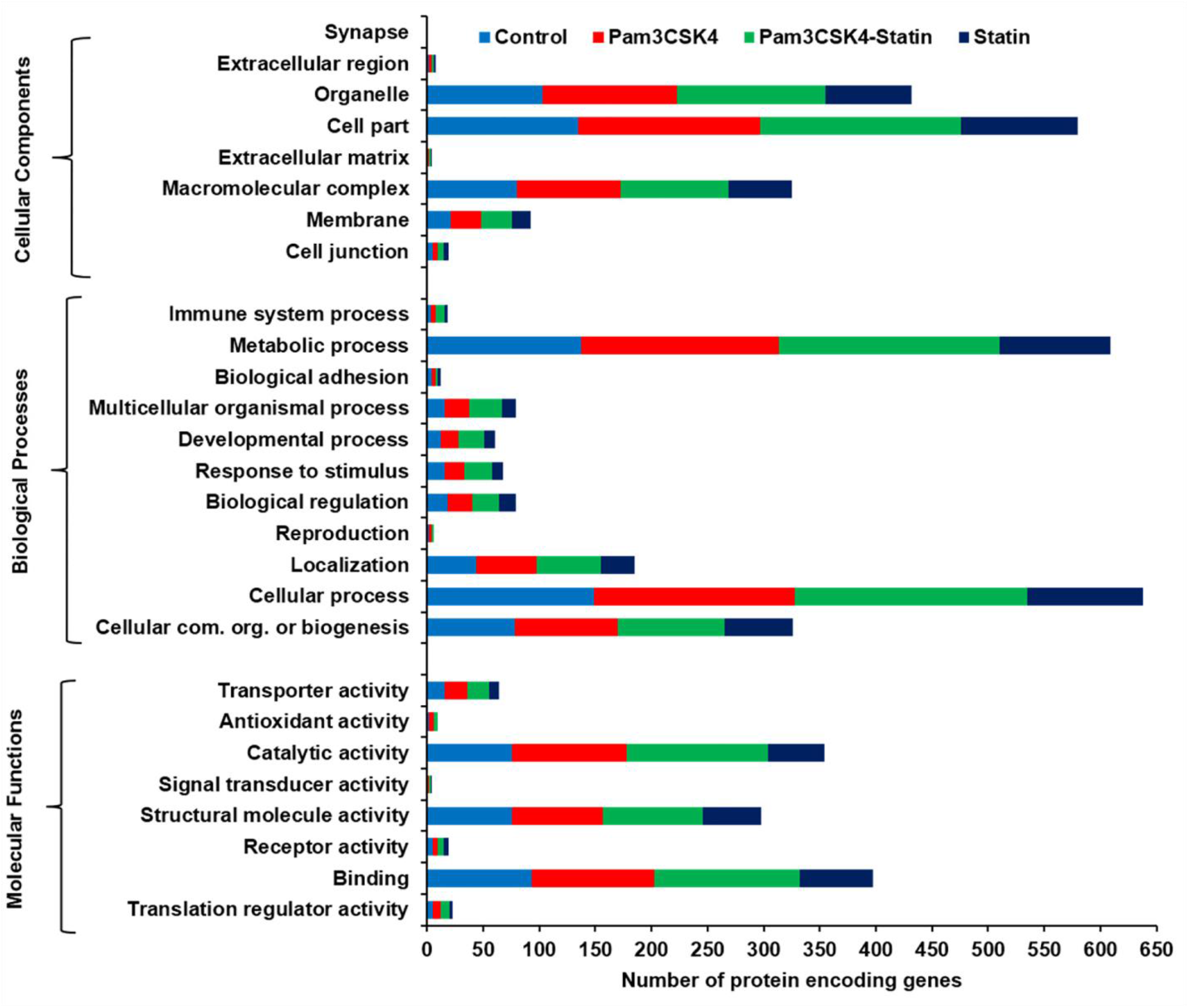
Gene ontology enrichment analysis. All identified and quantified proteins were subjected to gene ontology enrichment analysis based on molecular functions, biological processes, and cellular components.

### IPA-based TLR2-targeted protein interactions network

Core analysis was next performed by IPA (Ingenuity Systems) (37) in order to construct hypothetical interaction networks, canonical pathways, functions and disease pathways, and putative upstream regulators among the TLR2 interactome proteins identified. We found 25 classified networks according to top diseases and functions, within which a TLR2 protein-interacting network was ranked number eight (Table S5). The TLR2 protein network centered on cellular compromise, cellular function and maintenance, and cell death and survival. Given that we had by design targeted TLR2 in our interactome strategy, we next performed a TLR2-based network analysis (Fig. 5). Here, we found 32 interacting partner proteins. Expression of TLR2 itself was slightly increased by treatment with P3C, but sharply decreased during treatment with statin-P3C and statin (Figs. 5 and S3A-B). In this putative network, TLR2 interacted directly with heat shock protein 90-beta (HSP90B1) and the PI3-kinase p85 (PI3K p85) complex, and indirectly with vascular endothelial growth factor (VEGF). In the network, VEGF, in turn, is proposed to be regulated by interleukin enhancer-binding factor 3 (ILF3), which, in turn, is interconnected to heterogeneous nuclear ribonucleoprotein R (HNRNPD), far upstream element-binding protein 2 (KHSRP), ELAV-like protein 1 (ELAVL1), and ILF2. Of interest, ELVL1 protein is densely interconnected with 10 proteins experimentally observed in our pulldown. Furthermore, ELVL1 connects to PI3K family, HNRNPD, and KHSRP, highlighting the potential for complex intercommunications between proteins in the proposed TLR2 interactome (Fig. 5 and S3A-B). In this study, we found that ATP citrate lyase (ACLY) was highly increased (10.8-fold) upon treatment with P3C compared to statin-P3C and statin treatment (Figs. 5, S3A, and S3B). ACLY has recently been reported to support macrophage inflammatory activation (38), but has not previously been connected to TLR2 to our knowledge. Its suppression by statin treatment in our study interestingly identifies it as a putative mediator of the effect of statins on the inflammatory response.

**Figure 5.**
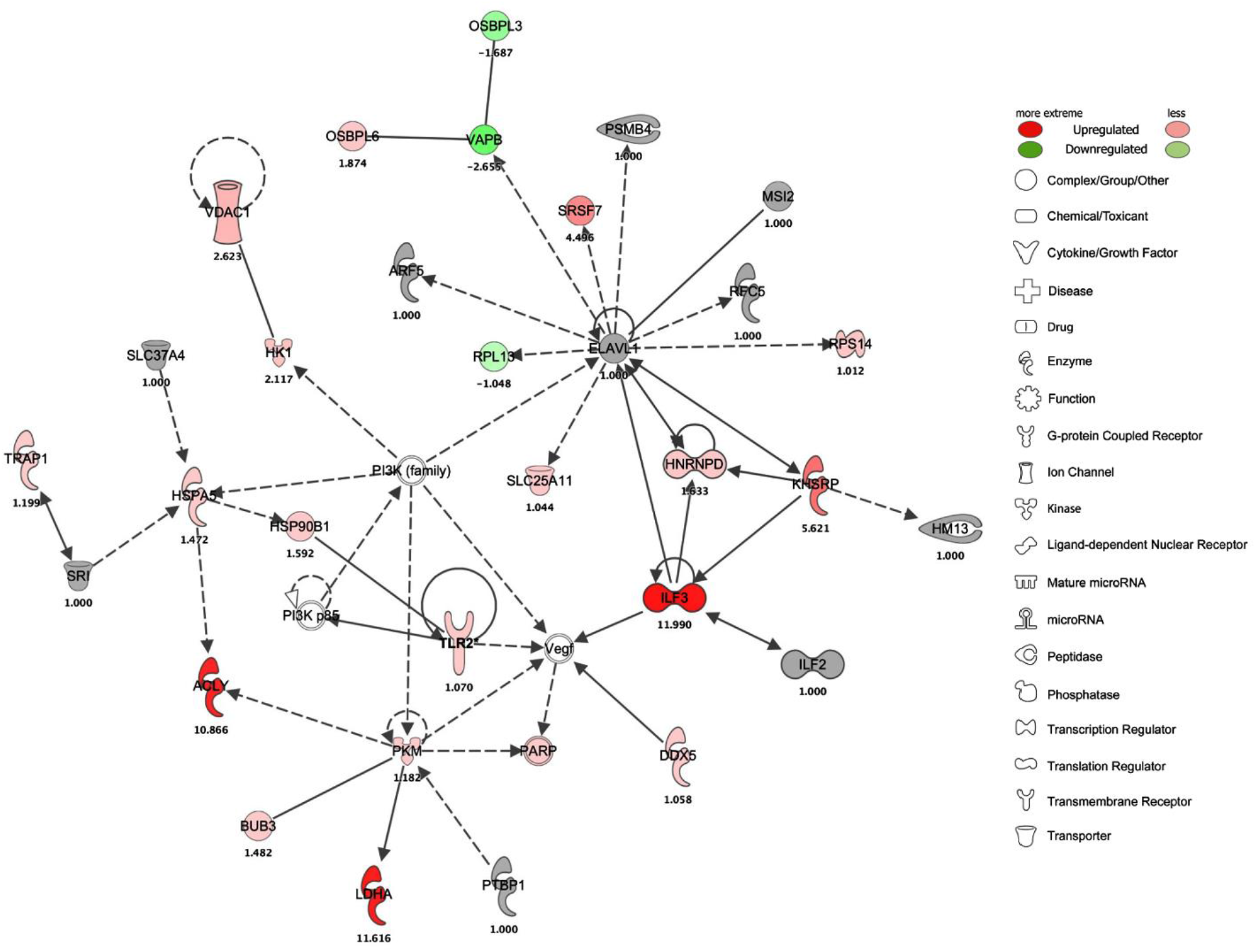
TLR2-targeted protein network with expression. The interaction network shown was generated using IPA bioinformatics software upon the treatment of Pam3CSK4 in HEK293 cells.

### Protein identification and interactions after cross-linking

In this study, we used two chemical crosslinkers, DUCCT and BS3, for covalent attachment of nearby proteins, aiming to improve recovery of low abundance and weakly interacting proteins. After DUCCT, we identified 605, 285, and 618 proteins under P3C, statin-P3C, and statin exposure conditions. After BS3, 365, 362, and 410 proteins were correspondingly identified under these three exposures (Table S2-S4). After stringent filtering among cross-linker control, treatment controls, and cross-linked samples, we exclusively identified 166 proteins in P3C-DUCCT, 47 proteins in statin-P3C-DUCCT, and 225 proteins in statin-DUCCT-treated samples (Figs. 6 and S4, Table S3). Correspondingly, we exclusively identified 32 proteins in P3C-BS3, 43 proteins in statin-P3C-BS3, and 40 proteins in statin-BS3-treated samples (Figs. S4-S5, Table S4). Consequently, considering total and exclusively identified proteins, DUCCT cross-linker enriched more TLR2-interacting proteins compared to BS3. After stringent filtering of the identified proteins among all exposure and crosslinking conditions individually and in combination, the data indicates that DUCCT exhibits superior efficiency to couple proteins across different treatment conditions compared to BS3 (Fig. S4). A protein-protein interaction network was constructed using the exclusively identified proteins due to DUCCT and BS3 treatments among the four cell exposure conditions (control, P3C, statin-P3C, statin), using the UniProt database via Cytoscape software (Figs. 6 and S5). A total of 325 DUCCT-exclusive proteins were used to generate the networks, containing 218 nodes and 320 edges (Fig. 6). As is evident, the highest node degree genes were RNA binding motif protein 8A (RBM8A; 35 edges), endoplasmic reticulum lipid raft-associated protein 2 (ERLIN2; 28 edges), eukaryotic translation initiation factor 4A3 (EIF4A3; 19 edges), RuvB like AAA ATPase 2 (RUVBL2; 16 edges), eukaryotic translation initiation factor 3 subunit B (EIF3B; 14 edges), splicing factor proline and glutamine rich (SFPQ; 14 edges), and transmembrane P24 trafficking protein 9 (TMED9; 13 edges). In parallel, 92 BS3-exclusive proteins were used to generate a network containing 121 nodes and 141 edges, within which G3BP stress granule assembly factor 1 (G3BP1) protein-coding gene showed high interaction with 3 node genes (Fig. S5).

**Figure 6.**
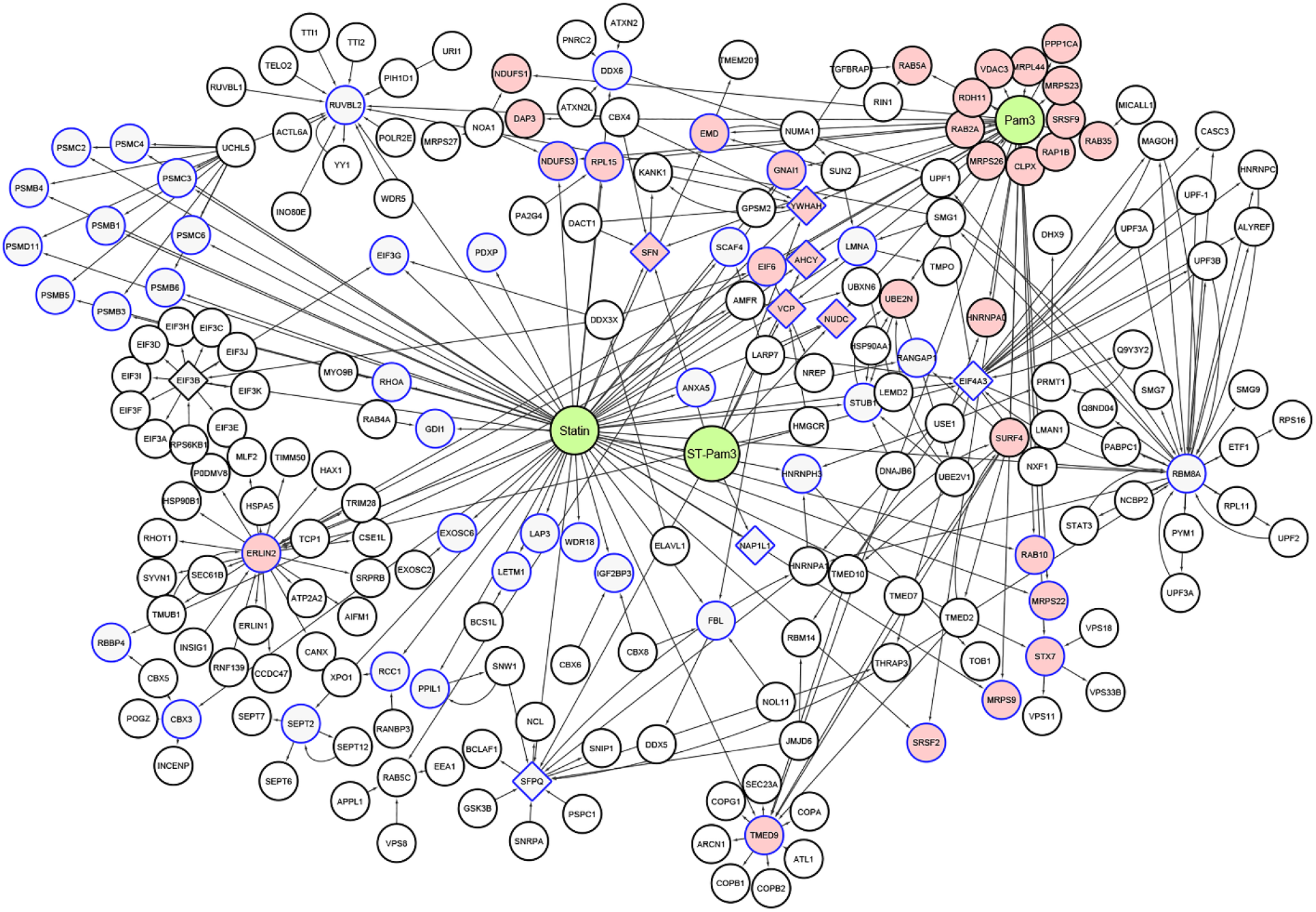
Protein interaction network of exclusively identified proteins by DUCCT crosslinking upon treatment with Pam3CSK4, statin-Pam3CSK4, and statin. Cytoscape (see methods section) was used to generate protein networks. The pink coloring indicates proteins identified in Pam3CSK4, diamond shapes indicate proteins identified in statin-Pam3CSK4 samples, and blue color (border color) indicates proteins identified in statin treated samples.

### Validation of selected proteins and their interacting partners

To verify the mass spectrometry-based protein data, we performed IP and immunoblot analysis on selected candidate proteins. Among the TLR2-interacting proteins identified, we focused our attention on alpha-centractin (ACTR1A) and myristoylated alanine-rich C kinase substrate-like protein 1 (MARCKSL1), based on their expression as well as their potential functions. In the HA-TLR2 interactome proteomics pulldown, ACTR1A was identified exclusively in the DUCCT-treated samples under the two exposure conditions of P3C and statin (Fig. 7A), whereas MARCKSL1 protein was detected only in statin-P3C and statin exposure conditions in the absence of crosslinker treatment (Fig. 7A), suggesting distinct patterns of responsiveness of these two proteins to P3C and statin. For validation, first, we performed immunoblot analysis of whole cell lysates to evaluate the expression status of these two proteins. Both ACTR1A and MARCKSL1 were highly upregulated in statin-P3C- and statin-treated samples compared to control and P3C-treated samples (Fig. 7B), suggesting that statins induce the expression of these two proteins in HEK293 cells. Next, HA-TLR2 IP samples were analyzed by immunoblot. We found that levels of ACTR1A co-precipitating with HA-TLR2 were significantly decreased in statin-treated cells (Fig. 7B). In order to further validate interactions of TLR2 with ACTR1A and MARCKSL1, we performed a reverse co-IP (i.e., immunoblot of TLR2 after ACTR1A IP). This revealed that TLR2 was highly increased in P3C- and statin-P3C-treated ACTR1A pull-down samples compared to control and statin-treated samples (Fig 7B). TLR2 was increased in P3C-, statin-P3C-, and statin-treated MARCKSL1 pull-down samples compared to control (Fig. 7B). Taken together, these findings suggest that P3C and statins enforce differential changes in the interaction of TLR2 with ACTR1A and MARCKSL1 in HEK293 cells. For further cross-validation, we performed immunocytochemistry on ACTR1A and TLR2 in the HEK293 cells. Here, we noted that in HEK293 cells TLR2 protein expression was inhibited by statin treatment, whereas ACTR1A protein was increased by statins (Fig. 8).

**Figure 7.**
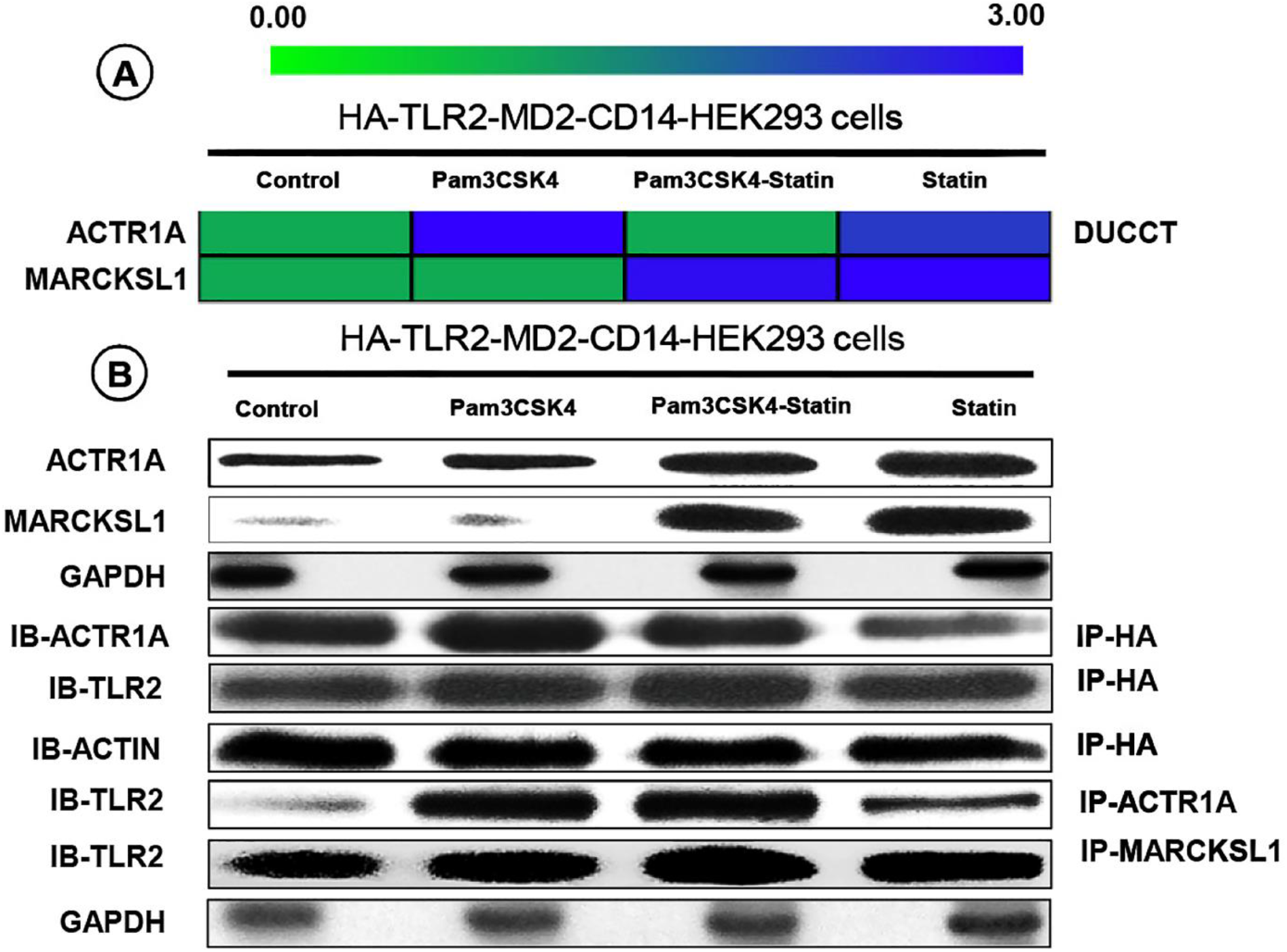
Validation of TLR2 protein interactors. (A) ACTR1A and MARCKSL1 proteins expression in HEK293 cells by LC-MS/MS. (B) ACTR1A and MARCKSL1 and their interactions were validated using immunoblotting (IB) and co-immunoprecipitation (IP) in HEK293 cells. All samples were treated with statin drug and bacterial ligand Pam3CSK4 except control.

**Figure 8.**
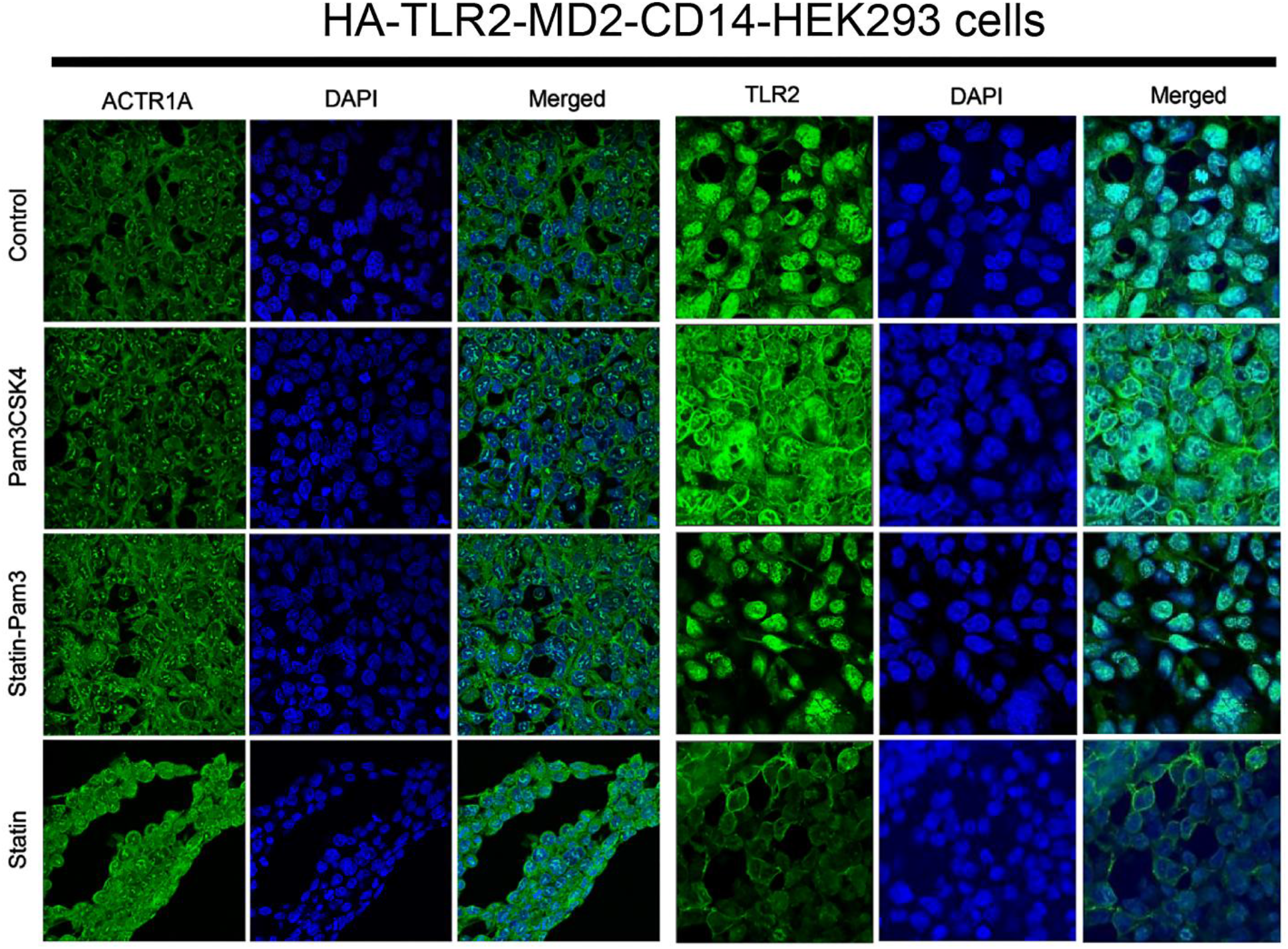
Cross-validation of candidate proteins. ACTR1A and TLR2 protein expressions were cross-validated using immunocytochemistry upon the treatment of Pam3CSK4 and statin in HEK293 cells.

### ACTR1A knockdown changes the levels of cytokines

To test for a possible function of ACTR1A in the TLR2 inflammatory response, we used siRNA to silence ACTR1A in HEK293 cells. After confirmation of siRNA efficiency in untreated cells (Fig. 9A), we analyzed expression of ACTR1A and of the pro-inflammatory genes tumor necrosis factor (TNFα), interleukin 6 (IL-6), and interleukin 8 (IL-8) in cells exposed to P3C, statin, and P3C-statin (Fig. 9). ACTR1A gene expression was successfully silenced by the siRNA under all treatment conditions (Fig. 9B). As expected, P3C induced robust TNFα (Fig. 9C). Of interest, statin treatment by itself did not change TNFα from control levels, but augmented the TNFα induction response to P3C. Whereas the TNFα response to P3C was not modified by silencing of ACTR1A, the TNFα response to combined P3C-statin treatment was significantly inhibited by ACTR1A silencing, suggesting that statins augment TLR2-dependent TNFα through a mechanism that requires ACTR1A. Under our experimental conditions, P3C did not induce IL-6 in HEK293 cells, although, interestingly, statin treatment itself modestly increased IL-6 (Fig. 9D). Finally, as with TNFα, statins modestly augmented P3C induction of IL-8. Induction of IL-8 by P3C and P3C-statin treatment were both significantly reduced by ACTR1A silencing, suggesting that ACTR1A is required for both of these TLR2 responses of the cell.

**Figure 9.**
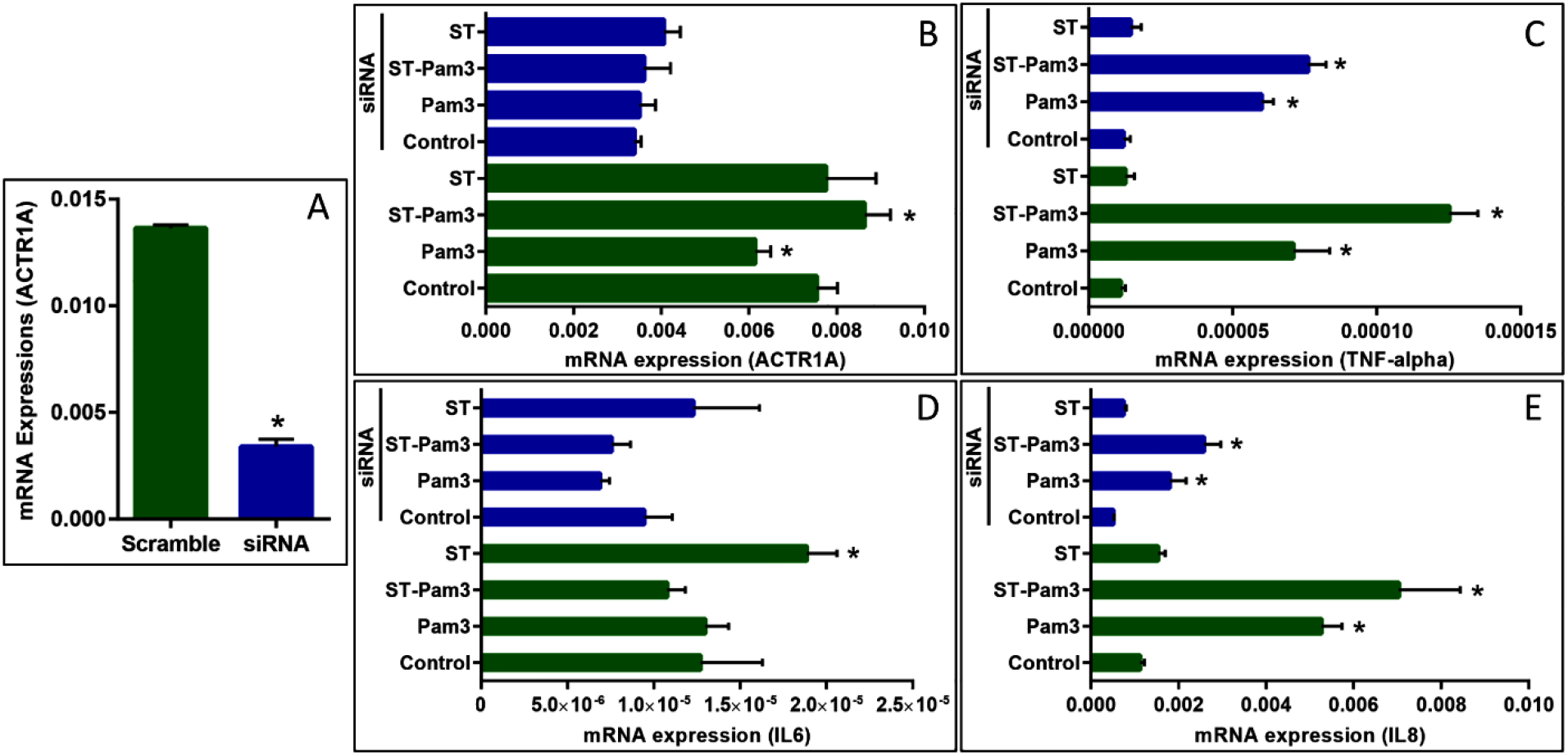
Silencing of ACTR1A modulates cytokine expression. (A) HEK293 cells were transfected with scramble siRNA and siRNA targeting ACTR1A, and then ACTR1A expression was analyzed by qRT-PCR. (B-D) After 48 hours with or without siRNA treatment, the relative mRNA expression of ACTR1A (B), TNF-alpha (C), IL6 (D), and IL8 (E) were analyzed by qRT-PCR upon treatment with statin and Pam3CSK4, as shown. All data are showed as mean ± SEM (n= 3 in each group) with *p<0.05.

## Discussion

Protein-protein interactions play a central role in signal transduction. Currently, IP coupled to high throughput mass spectrometry is popular for identifying target protein complexes (11, 39). Co-IP-based mass spectrometry has become a gold standard to study these interactions in a large-scale experimental design. Cross-linking before affinity or antibody pulldown aids in protein discovery, but has been hindered in many cases by the low abundance of target proteins and also by the analytic challenges of cross-linked peptides in large-scale proteomic studies. We have developed DUCCT, a dual cleavable crosslinker with a spacer chain distance of ~16 Å and a software package is currently under development to search the resulting cross-linked products.

In the present study, for the first time, we utilized two cross-linkers with different spacer chain lengths to study TLR2-interacting proteins. Aiming to test for possible effects of statins on the TLR2 interactome, we evaluated cells treated with the TLR2 ligand P3C, with statin, or with P3C following statin treatment. Novel interactors were identified through pulldown of TLR2 by a HA epitope tag, and further validated with biochemical approaches. Importantly, our data indicate that DUCCT enhances recovery of the TLR2 interactome and does so in a superior fashion to BS3. Of interest, gene ontology analysis of the interactors using PANTHER gene function classification showed nearly half or one-third of the identified proteins were involved in the protein binding category (33.96%) under molecular functions, cellular processes category (30.65%) under biological processes, and cell part category (39.69%) under cellular components (Fig. 4).

Computational biology techniques, in particular IPA and Cytoscape, were used to predict interaction partners and targeted protein-encoding genes following treatment of cells with P3C and/or statin. We predicted the TLR2-targeted proteins network in Fig. 5 using the identified proteins in the pull-down samples. In Table S5, IPA analysis predicted a TLR2 protein network involved in cell integrity including maintenance the cellular function, cell death and survival (40). Interestingly, IPA networks predicted direct interactions between TLR2 and HSP90B1, a protein detected by us in the TLR2 interactome (Fig. 5). HSP90B1 protein expression increased in P3C-stimulated samples, while decreasing in statin-P3C and statin-treated samples (Figs. 5 and S3A-B), suggesting that statin treatment may modify the proximal TLR2 interactome. The endoplasmic reticulum HSP90 isoforms are commonly involved in cell survival and proliferation (41). A previous study reported that HSP90B1 upregulation is associated with epithelial ovarian cancer and cancer cell survival (42). Moreover, bacterial infection and inflammation were increased in HSP90B1-deficient macrophages, which interfered with TLR signaling as well as function in the affected cells (43). Given this, we speculate that inhibition of HSP90B1 could be a potential strategy in cancer and anti-inflammatory therapy. Our interactome results further suggest that statins may be a potential tool for targeting HSP90B1 in the TLR2 cascade.

Protein-protein interaction network analysis upon treatment with P3C and statins along with DUCCT and BS3 revealed several proteins and protein complexes which are potential TLR2 interactors. RBM8A proteins were exclusively identified in statin-treated cells following DUCCT (Fig. 6). RBM8A proteins belong to the RNA-binding protein family. This family of proteins is believed to be involved in protein synthesis, cell cycle, apoptosis, and RNA splicing (44–47) and has also been shown to regulate intercellular immunity and production of inflammatory and anti-inflammatory cytokines (48). RBM8A is reported to be associated with the regulation of apoptosis, as inhibition of this protein increased cell apoptosis and reduced cell proliferations (45, 49). We speculate, based on our findings, that RBM8A may play a particularly important role in TLR2-mediated cell responses.

We validated two potential interacting partners of TLR2, namely ACTR1A and MARCKSL1, using immunoblot, reciprocal co-IP, and immunocytochemistry (Fig. 7–8). ACTR1A and MARCKSL1 proteins were remarkably upregulated by statin treatment in HEK293 cells (Fig. 7–8). The related protein MARCKS is known to be a regulator of actin cytoskeleton dynamics and MARCKSL1 protein is thought to share similar molecular functions. MARCKSL1 is expressed in a variety of different cells including astrocytes, fibroblasts and microglial cells and is upregulated in macrophages by LPS (50–53). MARCKS proteins crosslink actin filaments and maintain mechanical and structural properties of the actin cytoskeleton. During macrophage activation, MARCKS joins with actin or actin-binding proteins at sites of the cytoskeleton and plasma membrane (51, 54, 55). Both actin crosslinking and formation of actin bundles are inhibited by protein kinase C-dependent phosphorylation and binding of calmodulin to MARCKS proteins (55). MARCKS and MARCKSL1 proteins migrate from the cell membrane to cytosol upon phosphorylation or binding to calmodulin and thereby activate various signal transduction pathways by influencing actin dynamics and vesicular trafficking (56). Moreover, MARCKS proteins are involved in cell migration, adhesion, motility, membrane trafficking, mitogenesis, and spreading due to actin filament and microtubule reorganization (57, 58). Co-purification of MARCKSL1 with ACTR1A suggests their cooperation in actin cytoskeletal dynamics in the TLR2-mediated signaling response.

ACTR1A is cytoplasmic protein that is a part of the cytoskeleton, particularly microtubules. It appears as short filaments at the center of the dynactin complex that actively associate to dynein motor activity (59, 60). Cytoplasmic ACTR1A protein has also been found to play a vital role in actin filament assembly and organization (61) and intracellular vesicle trafficking. Prior studies have reported that macrophage activation by mycobacteria or cell wall lipoprotein p19 (TLR2 agonist) induces cytoskeletal rearrangement by TLR2-mediated phosphatidylinositol 3-kinase (PI3K) activation pathways (62). This activation of PI3K was necessary for the actin assembly and reorganization that underlies macrophage spreading and polarization (62). The actin cytoskeleton is also assembled in P3C-stimulated dendritic cells which enhances antigen generation and capture (63). Taken together, a key role for the actin cytoskeleton has been identified in TLR2-dependent immune responses, and our findings lead us to speculate that ACTR1A, as a novel TLR2 interactor, may play an important role in mediating this connection.

Interestingly, in our studies, ACTR1A was highly expressed following statin treatment and also co-purified with TLR2 in HEK293 cells, suggesting that it is a statin-sensitive TLR2 interactor (Fig. 7–8). Our co-IP-based mass spectrometry studies revealed that TLR2 interacts with ACTR1A in HEK293 cells upon statin and P3C treatment. The TLR2-ACTR1A interaction was confirmed with biochemical approaches. Further, to study the functional involvement of ACTR1A in TLR2 signaling pathways, we knocked down the expression of ACTR1A. Silencing of ACTR1A interestingly reduced pro-inflammatory cytokine expression in HEK293 cells, confirming an important role of ACTR1A in transducing the TLR2 pro-inflammatory signal. Future studies may be warranted to determine whether ACTR1A supports TLR2 signaling through linking TLR2 to the underlying cortical cytoskeleton.

Of note, we identified ACTR1A after crosslinking with DUCCT but not BS3. DUCCT has a larger cross-linker spacer chain compared to BS3 and may thus be superior for conjugating larger protein complexes and penetrating the plasma membrane. A part of large dynactin complex, ACTR1A function is not clearly known. This is the first time to our knowledge that ACTR1A and TLR2 interaction has been revealed by using co-IP cross-linking proteomics. MARCKS in the plasma membrane binds to dynamitin, a subunit of dynactin complex in cytoplasm (57). Co-purification of MARCKSL1 and ACTR1A following statin treatment suggests a role for cytoskeletal rearrangement in association with TLR2. Further functional studies are thus warranted to decipher the detailed mechanism of interactions between these newly discovered proteins in cytoskeletal dynamics and in statin-sensitive TLR2 signaling.

## Acknowledgments

We acknowledge funding from 1UA5GM113216-01, NIGMS, NIH to PI Dr. Chowdhury and also from the UT Proteomics Core Facility Network for a mass spectrometer. We also acknowledge start-up funding to Dr. Chowdhury from the University of Texas at Arlington. Dr. Michael Fessler is funded by the Intramural Research Program of the National Institute of Environmental Health Sciences, NIH (Z01 ES102005).

## Footnotes

### Author contributions

AHMK and SMC designed the experiment, JJA made and MF provided the HA-TLR2-HEK293 cell line, AHMK performed the experiment, AHMK and SMC analyzed the data, AHMK, MF and SMC wrote the manuscript, all authors read the manuscript.

## Abbreviations

HEK293T: HA-TLR2-MD2-CD14-HEK293 cells
XL: Cross-Linker
P3C: Pam3CSK4
ACTR1A: alpha-centractin
DUCCT: Dual Cleavable Cross-Linking Technology

