## supplementary information for "Crosslinking Proteomics Indicates Effects of Simvastatin on the TLR2 interactome and Reveals ACTR1A as a Novel Regulator of the TLR2 Signal Cascade"

Current location: [a] East Carolina Diabetes and Obesity Institute, Department of Physiology; East Carolina University, Greenville, NC 27834-4354, USA

### Supplementary Information

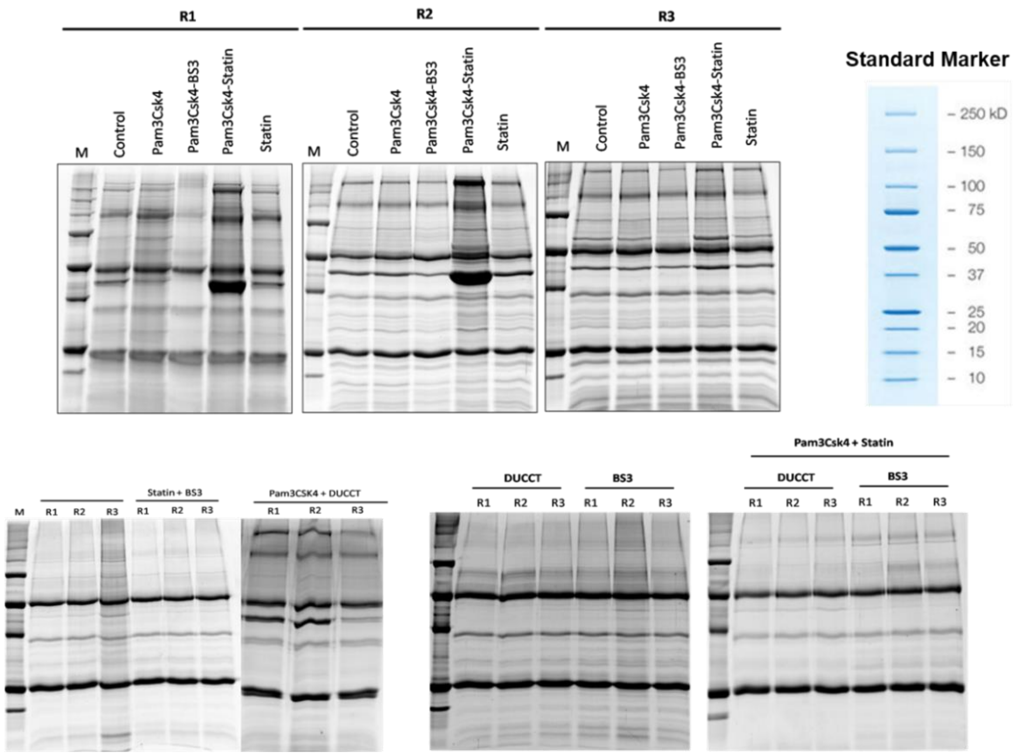


**Figure S1.** Representative gel images upon treatment with Pam3CSK4 and statin along with DUCCT and BS3 cross-linkers in HEK293 cells. Pull-down was performed using anti-HA magnetic beads, and then samples were separated on SDS-PAGE gels. SDS-PAGE gels of three biological replicates stained with Sypro Ruby are presented.

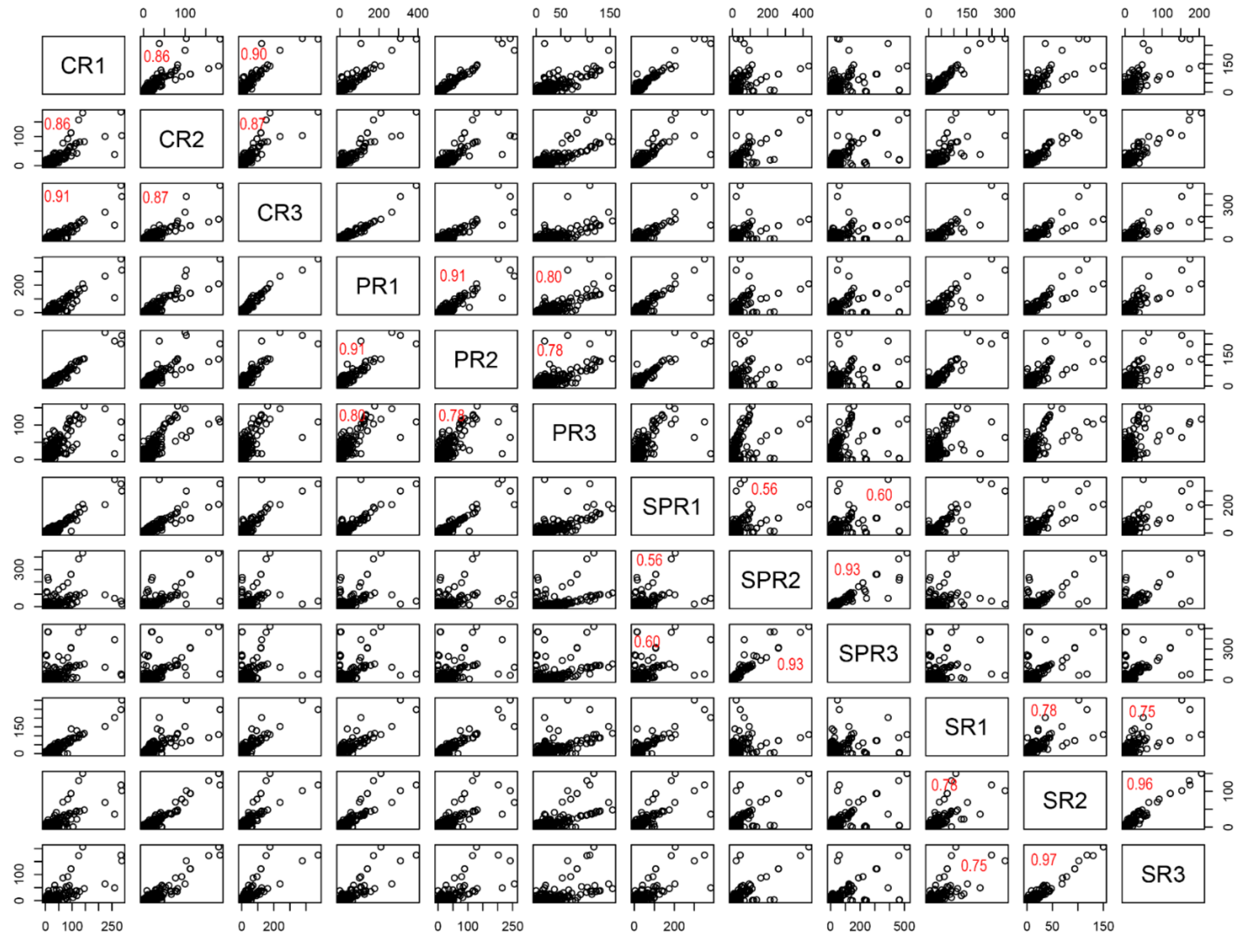


**Figure S2.** Scatter plots and pair wise correlations show significant correlation patterns among the replicate samples with treatments Pam3CSK4 and statin in HEK293 cells. The PSMs (spectral counts) of proteins of replicate samples are plotted against each protein on the *x*-axis and *y*-axis, correspondingly. Every spot symbolizes the abundance of a protein and corresponds to Pearson's correlation coefficient ($R^2$) of 1. The scatter plot and pair wise graph was generated by R package ver. 3.3.1. Indications: C, Control; P, Pam3CSK4; SP, statin-Pam3CSK4; and S, statin

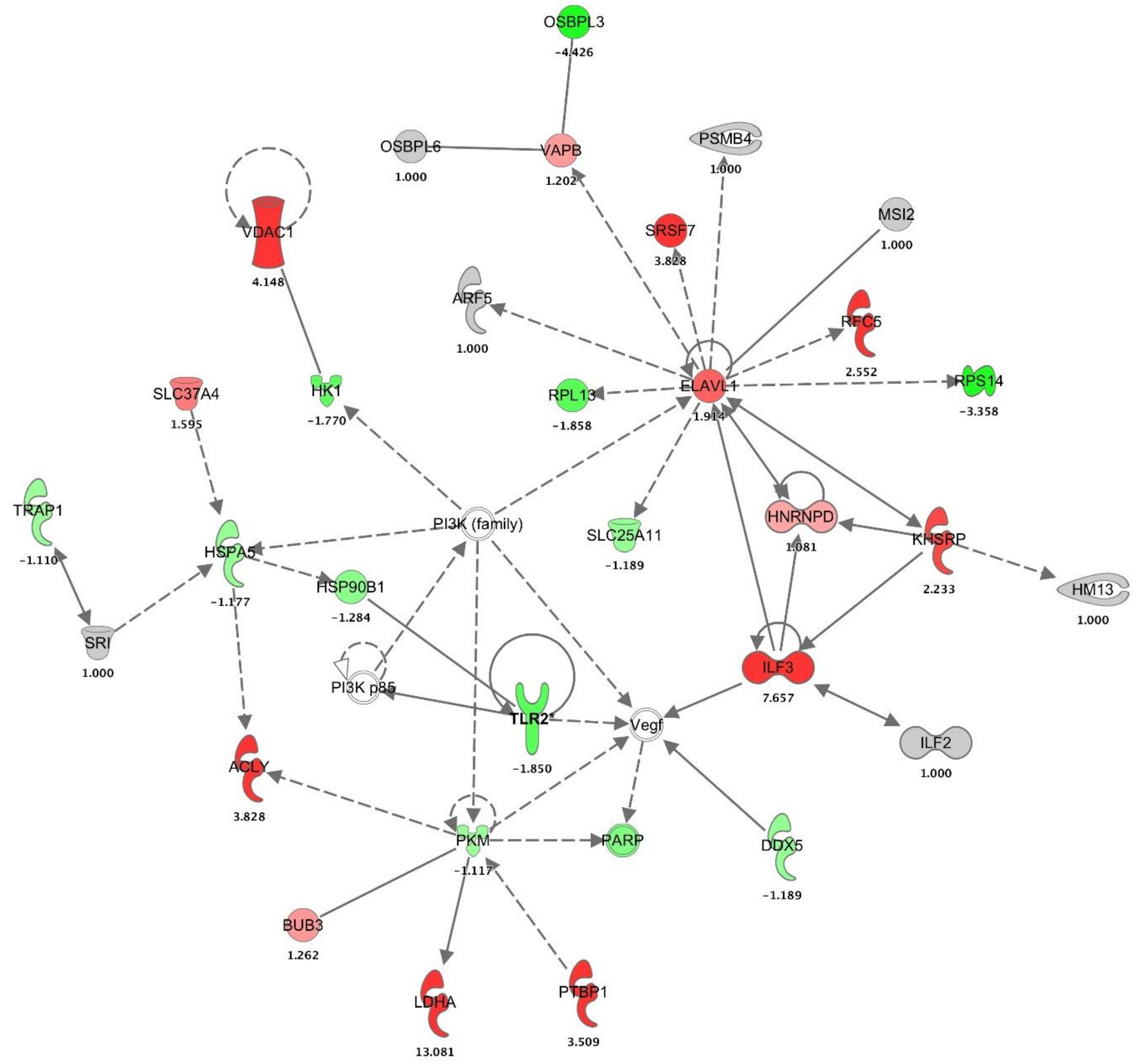


**Figure S3A.** TLR2-targeted protein network with expression. The interaction network was generated using IPA bioinformatics software upon the treatment of statin-Pam3CSK4 in HEK293 cells. Based on top diseases and functions, this network involves cellular compromise, cellular function and maintenance, cell death and survival.

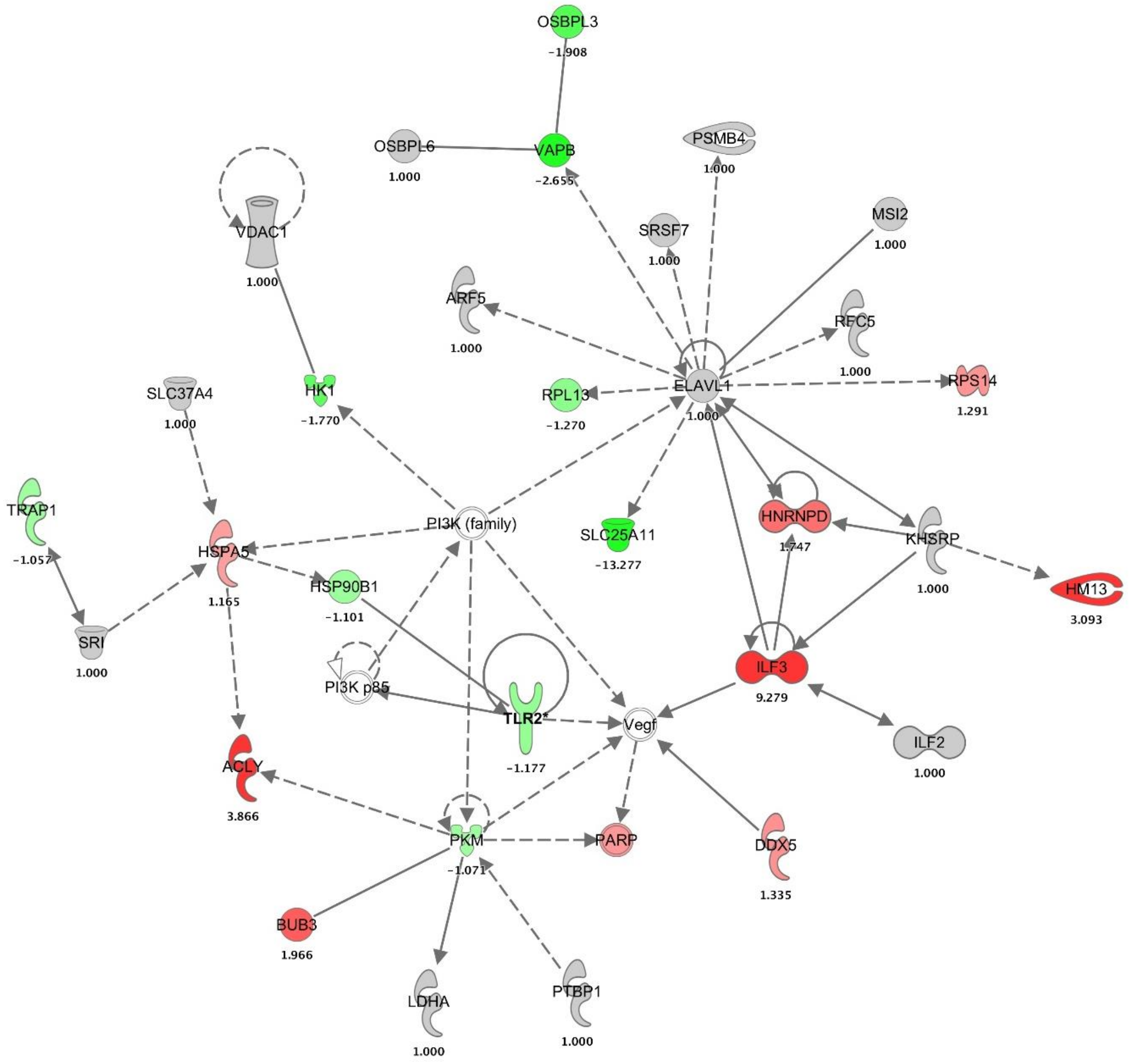


**Figure S3B.** TLR2-targeted protein network with expression. The interaction network was generated using IPA bioinformatics software upon treatment of statin in HEK293 cells. Based on top diseases and functions, this network involves cellular compromise, cellular function and maintenance, cell death and survival.

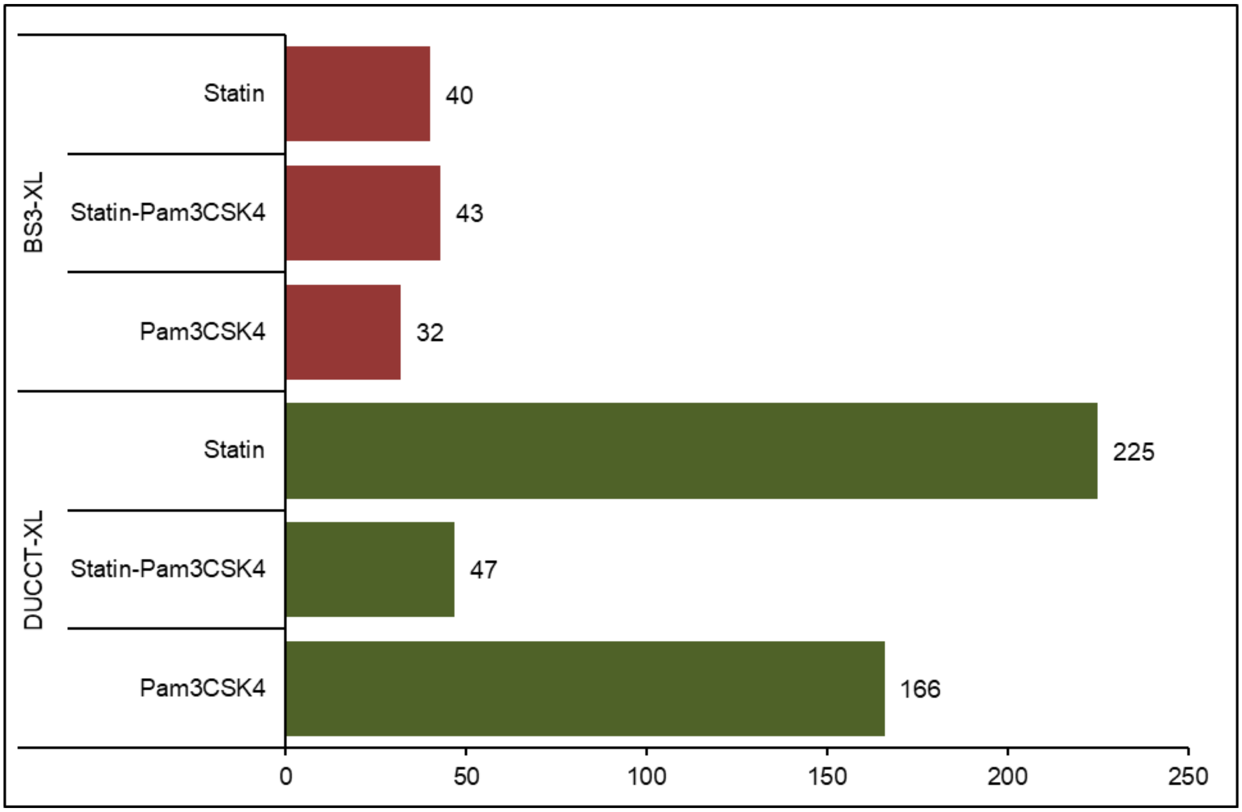


**Figure S4.** The graph shows the protein exclusively identified upon treatment with Pam3CSK4 and statin along with DUCCT and BS3 cross-linkers. Those proteins were filtered rigorously among control, crosslinker-control and treated samples with or without cross-linkers.

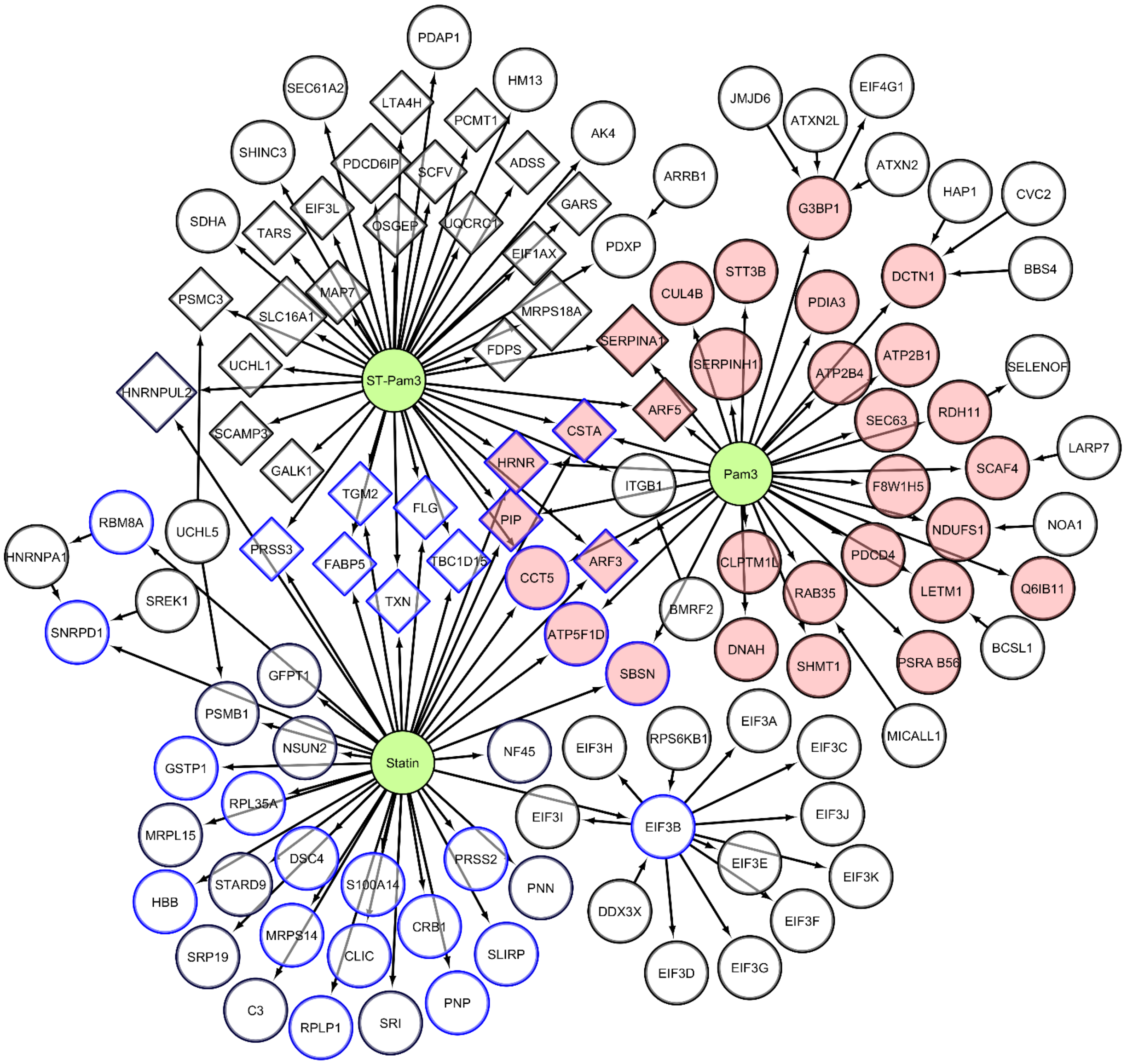


**Figure S5.** Protein interaction network of proteins exclusively identified by BS3 crosslinker upon treatment with Pam3CSK4, statin-Pam3CSK4, and statin. Cytoscape (see methods section) was used to generate the protein networks. The pink colors (fill color) indicate Pam3CSK4, diamond shapes indicate statin-Pam3CSK4, and blue color (border color) indicates statin conditions.

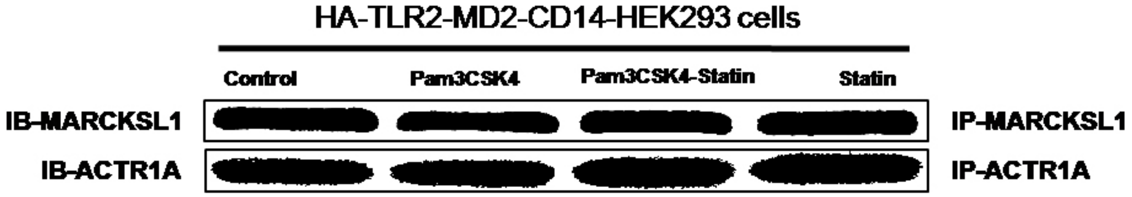


**Figure S6.** Expressions of ACTR1A and MARCKSL1 proteins analyzed by Western blot analysis for confirming the co-IP efficiency. The pulldown was performed through MARCKSL1 and ACTR1A antibody from the samples treated with P3C and statin-P3C in HEK293 cells. Then, membranes were immunoblotted with MARCKSL1 and ACTR1A antibody. The figure shows confirmation and efficiency of co-IP with those antibodies.
