## Supplementary material for "Crosslinking Proteomics Indicates Effects of Simvastatin on the TLR2 interactome and Reveals ACTR1A as a Novel Regulator of the TLR2 Signal Cascade": Table S1

List of primers for qRT-PCR analysis

| **Genes** | **Species** | **Forward Sequences** | **Reverse Sequences** |
| --- | --- | --- | --- |
| ACTR1A | Human | 5՛-CAAGCACGTTCGTGTCATGGCA-3՛ | 5՛-CGTTCCAATCCTTGACGATGCC-3՛ |
| TNF-α | Human | 5՛-CTCTTCTGCCTGCTGCACTTTG-3՛ | 5՛-ATGGGCTACAGGCTTGTCACTC-3՛ |
| IL-6 | Human | 5՛-AGACAGCCACTCACCTCTTCAG-3՛ | 5՛-TTCTGCCAGTGCCTCTTTGCTG-3՛ |
| IL-8 | Human | 5՛-GAGAGTGATTGAGAGTGGACCAC-3՛ | 5՛-CACAACCCTCTGCACCCAGTTT-3՛ |
| GAPDH | Human | 5՛-GTCTCCTCTGACTTCAACAGCG-3՛ | 5՛-ACCACCCTGTTGCTGTAGCCAA-3՛ |
